# Biobank of genetically defined murine prostate cancer tumoroids uncovers oncogenic pathways and drug vulnerabilities driven by PTEN-loss

**DOI:** 10.1101/2025.03.14.643296

**Authors:** Jessica Kalla, Thomas Dillinger, Zlata Pavlovicova, Reema Jacob, Emine Atas, Anil Baskan, Kristina Draganić, Andreas Tiefenbacher, Tanja Limberger, Theresia Mair, Gabriel Wasinger, Ludovica Villanti, Stefan Kubicek, Lukas Kenner, Gerda Egger

## Abstract

Prostate cancer (PCa) is the second most common cancer in men and shows high inter- and intra-patient heterogeneity. Consequently, treatment options are limited and there is a lack of representative preclinical models. Here we establish a comprehensive biobank of murine organoids and tumoroids that reflect common patient mutations. We demonstrate that the deletion of *Pten* alone, or in combination with *Stat3*, or *Tp53*, drives the activation of cancer-related pathways in both prostate organoids and tumour-derived tumoroids. A medium-throughput drug screen identified two potent compounds, the PDPK1/AKT/FLT dual pathway inhibitor and the sirtuin inhibitor tenovin-6, which effectively suppressed tumoroid proliferation. Notably, these compounds also inhibited the growth of several human PCa cell lines and displayed synergistic effects when combined with the standard-of-care antiandrogen enzalutamide. Together, our findings provide evidence that murine tumoroids are versatile preclinical models for studying PCa tumorigenesis and drug sensitivities to develop novel therapeutic options for PCa patients.

## Introduction

Prostate cancer (PCa), the second leading cause of cancer-related death in men worldwide^1^, is characterised by a diverse mutational landscape and high inter- and intra-patient heterogeneity.^2–4^ The malignant transformation of the normal prostate gland, which consists of luminal and basal epithelial cells, to PCa is a multifactorial process. Different driver events lead to the development of adenocarcinoma lesions that ultimately progress to metastatic disease.^5,6^ Radical prostatectomy, radiation therapy, and subsequent androgen deprivation therapy represent the primary treatment options for localised disease.^7^ However, despite initial response to these therapies, many patients eventually develop castration-resistant PCa and metastases, posing a significant therapeutic challenge due to limited treatment options and poor prognosis.^8^

Even though PCa is a very heterogeneous disease, some mutational patterns can be found in a large subgroup of patients.^9^ The loss of the tumour suppressor *PTEN* and thus an activation of the PI3K/AKT pathway, is one of the most common mutations found in PCa with an incidence of ∼20% in primary cases, and ∼50% in metastatic disease.^10^ *TP53* is also commonly mutated in PCa patients, and mutations in *TP53* frequently occur in combination with *PTEN* deletions.^9,11^ As *STAT3* is upregulated in many cancer types including PCa^12,13^, the inhibition of the IL-6/STAT3 signalling axis has been reported as a therapy approach for PCa.^14^ However, in mice the loss of *Stat3* in combination with the loss of *Pten*, a co-deletion observed in 66% of patients, led to a more aggressive and invasive phenotype, highlighting the dual role of STAT3 for PCa.^15–17^

Despite significant advances in our understanding of this disease, the development of effective therapies has been hindered by the lack of robust preclinical models that recapitulate the complex biology and treatment response of PCa patients.^18^ Only a hand-full of 2D cell lines derived from the human healthy prostate, or primary, and metastatic PCa exist. Since these cell lines consist of only one cell type, they do not fully recapitulate the *in vivo* tissue function and signalling of PCa tumours.^19^ Organoids and tumoroids, 3D *in vitro* models derived from primary patient tissue or animal samples, have emerged as a promising platform for cancer research.^20^ Even though it is possible to generate human prostate organoids and PCa tumoroids^21–24^, it is not possible to maintain these models in culture for extended passages. Primary PCa tumoroids get overgrown by healthy cells and most 3D models stop proliferating due to suboptimal medium and matrix conditions.^25–29^ Thus, PCa research has mainly relied on murine organoids and tumoroids, which can be maintained *in vitro* indefinitely, and provide a novel tool to study PCa tumorigenesis and therapy response.^23,25,30–32^

As the influence of different genetic mutations of murine prostate organoid and PCa tumoroid models on gene expression and drug response has not been studied extensively, we focused on establishing a biobank of organoids and tumoroids derived from wildtype (WT) or transgenic mice, respectively. Additionally, we generated PCa tumoroids by inducing the deletion of *Pten*, *Stat3*, and *Tp53* in WT organoids *in vitro*. Interestingly, the deletion of the target genes induced the deregulation of metabolic pathways in all knockout (KO) tumoroids. In addition, a medium-throughput compound screen identified the PDPK1/AKT/FLT dual pathway inhibitor (DPI) (also called KP372-1)^33–35^ and the epigenetic modifier tenovin-6 (T6), a sirtuin inhibitor and TP53 activator^36^, as promising agents. These compounds effectively inhibited the growth of murine tumoroid models and several human PCa cell lines. Thus, murine tumoroids provide reliable preclinical models for PCa research and could be used to identify additional novel treatments for PCa patients based on their genetic background.

## Results

### Establishment and genetic stability of murine PCa tumoroids reflecting patient mutations

To highlight the importance of modelling genetic mutations of *PTEN*, *STAT3*, and *TP53*, we analysed publicly available RNA sequencing data from primary PCa patients of the PRAD-TCGA dataset.^37^ A lower expression of the tumour suppressor *PTEN*, or the transcription factor *STAT3*, significantly correlated with shorter overall survival time (Figure 1A). Additionally, patients carrying mutations in the tumour suppressor *TP53* had a significantly shorter survival time compared to patients with a WT *TP53* gene. Together, this data underlined the important role of these genes for PCa tumorigenesis and patient prognosis.

**Figure 1:**
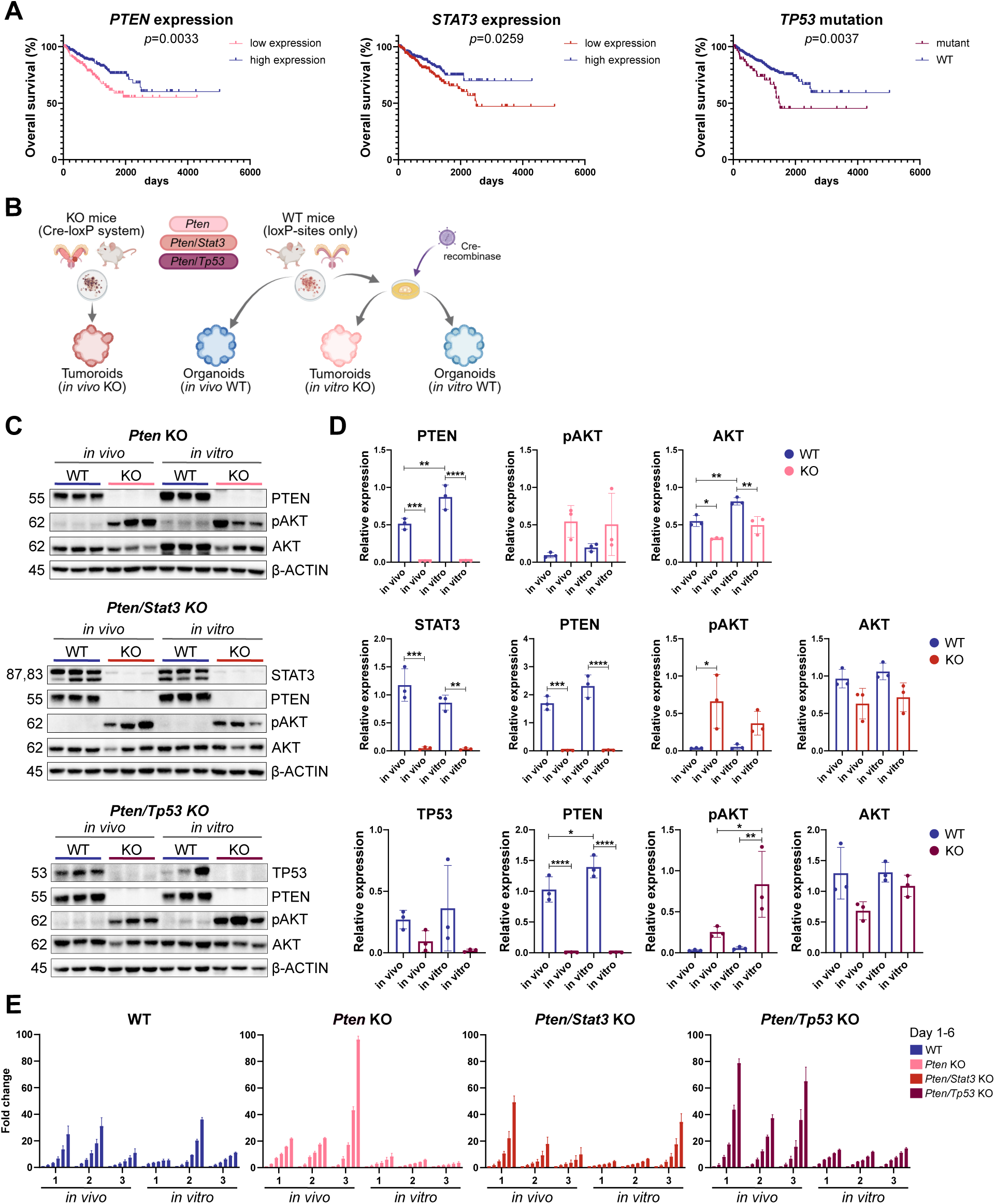
Establishment and genetic stability of murine PCa tumoroids reflecting patient mutations. **A.** Kaplan-Meier survival analysis of human PCa patients based on TCGA-PRAD RNA sequencing data for expression of *PTEN* (left), and *STAT3* (middle), or whole-genome/exome sequencing for *TP53* mutation status (right).^93^ Statistical analysis was done using GraphPad Prism 8.0.2 (Mantel-Cox test). **B**. Overview of the experimental design of this study. Tumoroids were derived from prostate tumours of mice with a genetic deletion based on the Cre-loxP system of *Pten* alone, or in combination with *Stat3*, or *Tp53* (*in vivo* KO). Healthy organoids were derived from Cre-negative mice with loxP-sites for the genes of interest (*in vivo* WT). Organoids with loxP-sites were then either transduced with a functional Cre-recombinase to induce the deletion of the genes (*in vitro* KO) or a non-functional Cre-recombinase as a control (*in vitro* WT). **C**. Western blot analysis of murine *in vivo* and *in vitro* organoids and KO tumoroids for indicated genotypes for PTEN, STAT3, TP53, phospho-AKT (pAKT), total AKT (AKT), and representative β-ACTIN as loading control. All samples shown in the *Pten/Tp53* KO blot were treated with CoCl_2_ to induce TP53 expression. **D**. Quantification of protein expression of Western blots shown in (C) (top: *Pten* KO, middle *Pten/Stat3* KO, bottom: *Pten/Tp53* KO). Bar graphs represent relative band intensity of proteins of interest normalised to β-ACTIN as loading control. Data are presented as means of triplicates ± SD. Statistical analysis was performed using Image Lab 6.1 and GraphPad Prism 8.0.2 (One-way ANOVA, Tukeýs test). *p* > 0.05 if not specified otherwise, \**p* ≤ 0.05; \*\**p* ≤ 0.01, \*\*\**p* ≤ 0.001, **** *p* ≤ 0.0001. **E**. Bar graphs depicting proliferation rates of *in vivo* and *in vitro* organoids and tumoroids (1-3 = biological replicates/single clones) for all genotypes over six days normalised to day 1 (fold change). Of note, *in vitro* organoid line WT1 has the same maternal line as *in vitro Pten* KO 1-3, WT2 corresponds to *in vitro Pten/Stat3* KO 1-3, and WT3 corresponds to *in vitro Pten/Tp53* KO 1-3. Each bar represents technical triplicates ± SD per organoid/tumoroid line.

To develop reliable preclinical models for PCa research, we took advantage of previously established conditional murine PCa models that harbour deletions of genes highly relevant for human PCa, including *Pten* single KO^38^, *Pten/Stat3*^16^, or *Pten*/*Tp53* double KO (dKO)^39^ (Figure 1B). Tumoroids were derived from the tumours of these mice at 19 weeks of age^40^ (*in vivo* KO), while WT organoids were generated from healthy prostates of Cre-negative mice carrying loxP-sites for the respective genes (*in vivo* WT). These WT organoids were subsequently transduced with a tamoxifen-inducible Cre-recombinase to induce the deletion of the respective genes (*in vitro* KO), to investigate the effects of these mutations on malignant transformation. In addition, *in vivo* WT organoids were transduced with a non-functional Cre-recombinase as a control (*in vitro* WT). Taken together, we generated an extensive biobank of murine organoids and tumoroids reflecting common PCa patient mutations associated with different stages of tumour aggressiveness.

The stable KO of the genes of interest in the *in vivo* and *in vitro* tumoroid models was confirmed on DNA and RNA level, whereby the deletion of the targeted exons for *Pten*, *Stat3*, and *Tp53* on DNA level (Figure S1A,B) resulted in the complete loss of gene expression in the KO tumoroids (Figure S1C). While all healthy organoid lines showed an expression of PTEN on protein level, the loss of PTEN and a subsequent activation of the PI3K/AKT pathway was seen in the KO tumoroids (Figure 1C,D). In addition, the absence of the STAT3 protein was confirmed in the *Pten/Stat3* dKO tumoroids. To visualise the expression or loss of TP53, we treated all organoid and tumoroid lines with cobalt chloride (CoCl_2_) leading to increased stability of TP53.^41^ While a heterogeneous expression of TP53 was seen in the WT organoids, the protein was lost completely in the *Pten/Tp53* dKO tumoroids (Figure 1C,D, bottom). In summary, all 3D models showed a clear loss of the respective proteins of interest, and an activation of the pro-tumorigenic PI3K/AKT pathway. Importantly, all *in vitro* organoids and KO tumoroids stably reflected the protein expression levels of their *in vivo* counterparts.

Based on the negative influence of mutations in *PTEN*, *STAT3*, and *TP53* on PCa patient survival, and the activation of the PI3K/AKT pathway promoting proliferation^42^, we expected a growth advantage of the KO tumoroids compared to WT organoids. Interestingly, we observed heterogenous proliferation rates among WT organoids and KO tumoroids, with *Pten/Tp53* dKO tumoroids showing the highest proliferation rate on average (Figure 1E). As organoid growth medium was optimised for the growth of healthy cells, we hypothesise that the proliferation rate mainly depends on the medium composition.^21^ Overall, *in vivo* KO tumoroids exhibited a noticeable trend of accelerated proliferation compared to WT organoids. In addition, exponential growth patterns were observed primarily in *in vivo* KO tumoroids, while *in vitro* KO tumoroids mainly exhibited linear growth patterns.

### Murine organoids and PCa tumoroids reflect the morphology of their tissue of origin

To investigate whether the organoids and tumoroids with different genetic backgrounds stably reflect their tissue of origin, we performed histo-morphological analyses including immunohistochemistry (IHC) on tissues and corresponding 3D models (Figure 2, Figure S1D). The murine healthy prostate tissue consists of glands made up of a two-layered epithelium, visible in haematoxylin and eosin (HE) staining and IHC for CK8-positive luminal cells and fewer P63-positive basal cells (Figure 2A). While there are nearly no proliferating cells expressing KI67 in the WT tissue, complex multi-layered, partly cribriform, and invasive glands with increased KI67 expression were observed in the KO tumours. KO tissues were also characterised by an increase in CK8-positive invasive cells and scattered basal cells.

**Figure 2:**
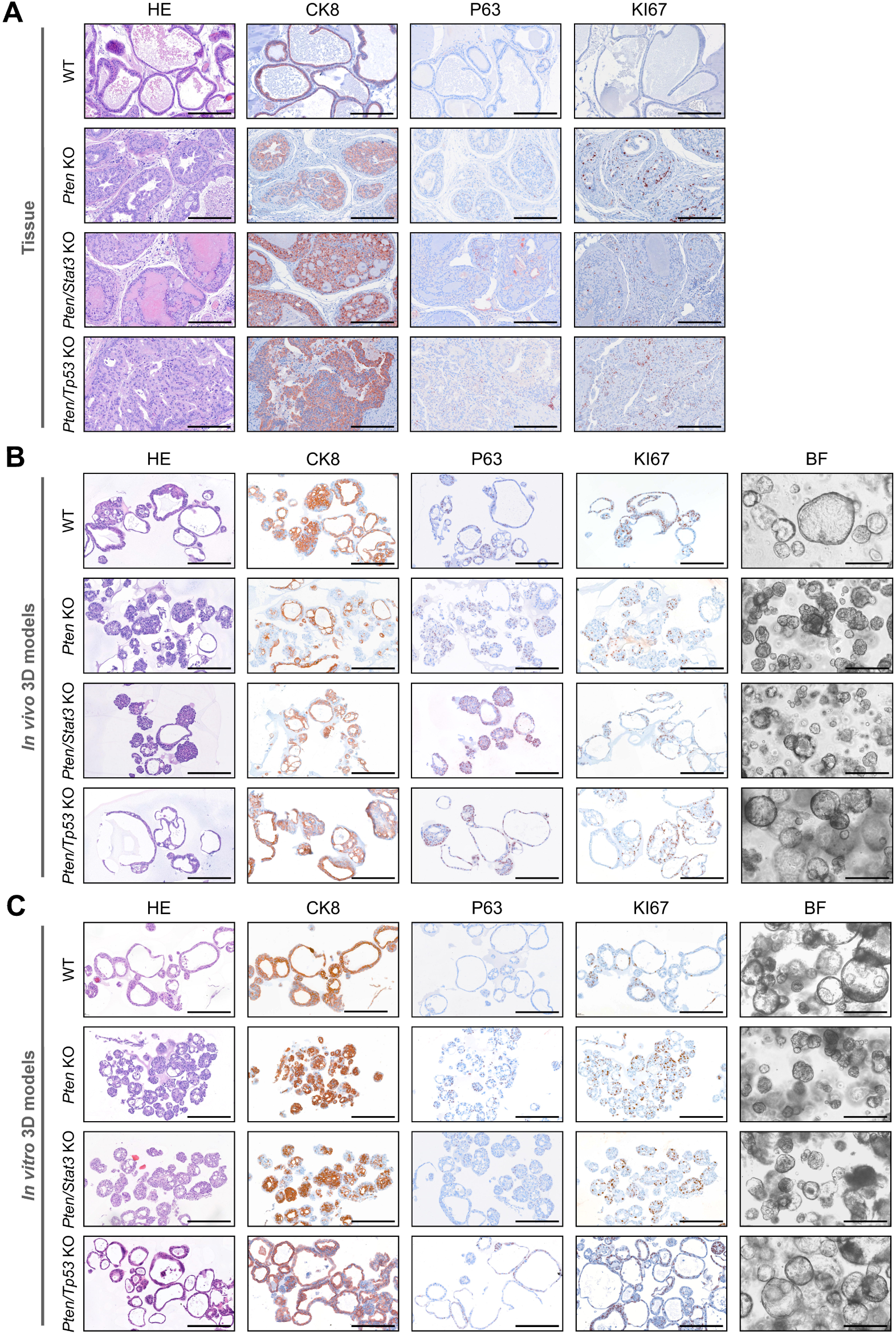
Murine organoids and PCa tumoroids reflect the morphology of their tissue of origin. **A**. Comparison between WT prostate tissues and prostate tumours of the *Pten* KO, *Pten/Stat3* KO, and *Pten/Tp53* KO PCa mouse models stained with haematoxylin and eosin (HE), or antibodies against KI67 (proliferation), CK8 (luminal cell marker), or P63 (basal cell marker). One representative mouse per genotype is shown (N=2). See also Figure S1D. Scale bar 200µm. **B**. Comparison between *in vivo* WT organoids and KO tumoroids for all genotypes. Organoids/tumoroids derived from murine tissues depicted in (A) are shown. In addition to HE, KI67, CK8, and P63 immunohistochemistry (IHC) stainings, bright-field (BF) microscopic images of 3D models are shown. Scale bar 200µm. **C**. Comparison between *in vitro* WT organoids and KO tumoroids for all genotypes. In addition to HE, KI67, CK8, and P63 IHC stainings, bright-field (BF) microscopic images of 3D models are shown. One representative line per genotype is shown (N=3). See also Figure S1D. Scale bar 200µm.

The *in vivo* WT organoids formed mostly large and hollow structures reflecting normal prostate glands, while the *Pten* KO and *Pten/Stat3* dKO tumoroids displayed a compact tumoroids displayed a slightly different growth pattern and formed the largest tumoroid spheres among all 3D models. In line with previous literature^23,43^, we mostly observed organoids and tumoroids consisting of both luminal and basal cells, with few structures consisting of only one cell type. The proliferation marker KI67 was expressed at similar levels in all organoid and tumoroid lines. Importantly, the *in vitro* deletion of *Pten* alone, or in combination with *Stat3*, or *Tp53* in WT organoids resulted in morphological changes reflected by dense growth patterns as previously observed in *in vivo* KO tumoroids (Figure 2C). Moreover, all *in vitro* KO tumoroids showed cancer-specific increased nuclear atypia in comparison to their healthy counterparts. CK8/P63 distribution and proliferation marked by KI67 were comparable to WT organoids and *in vivo* KO tumoroids. In conclusion, the phenotypic changes in organoid morphology possibly indicate the malignant transformation of WT organoids after genetic deletion of the target genes.

### Transcriptomic analysis of PCa tumoroids reveals upregulation of oncogenic signalling and alterations in metabolic pathways

To investigate the impact of PCa-specific mutations on gene expression and signalling, we performed bulk RNA sequencing on *in vivo* WT and KO 3D models. Principal component analysis of biological replicates showed heterogeneity both among and within the different genotypes, with two out of three biological replicates clustering closely together (Figure 3A). The most considerable heterogeneity was apparent in the *Pten/Tp53* dKO tumoroids. Unsupervised hierarchical clustering of the top 1000 most variably expressed genes reflected the previously observed heterogeneity of organoid and tumoroid lines (Figure 3B). Interestingly, *in vivo* 3D models separated independent of their genotype into two main groups, both containing organoids and tumoroids. Differential gene expression between these two groups identified a deregulation of genes and pathways involved in cell cycle regulation and mitosis (Figure S2A-E), highlighting the major impact of different proliferation rates on overall gene expression.

**Figure 3:**
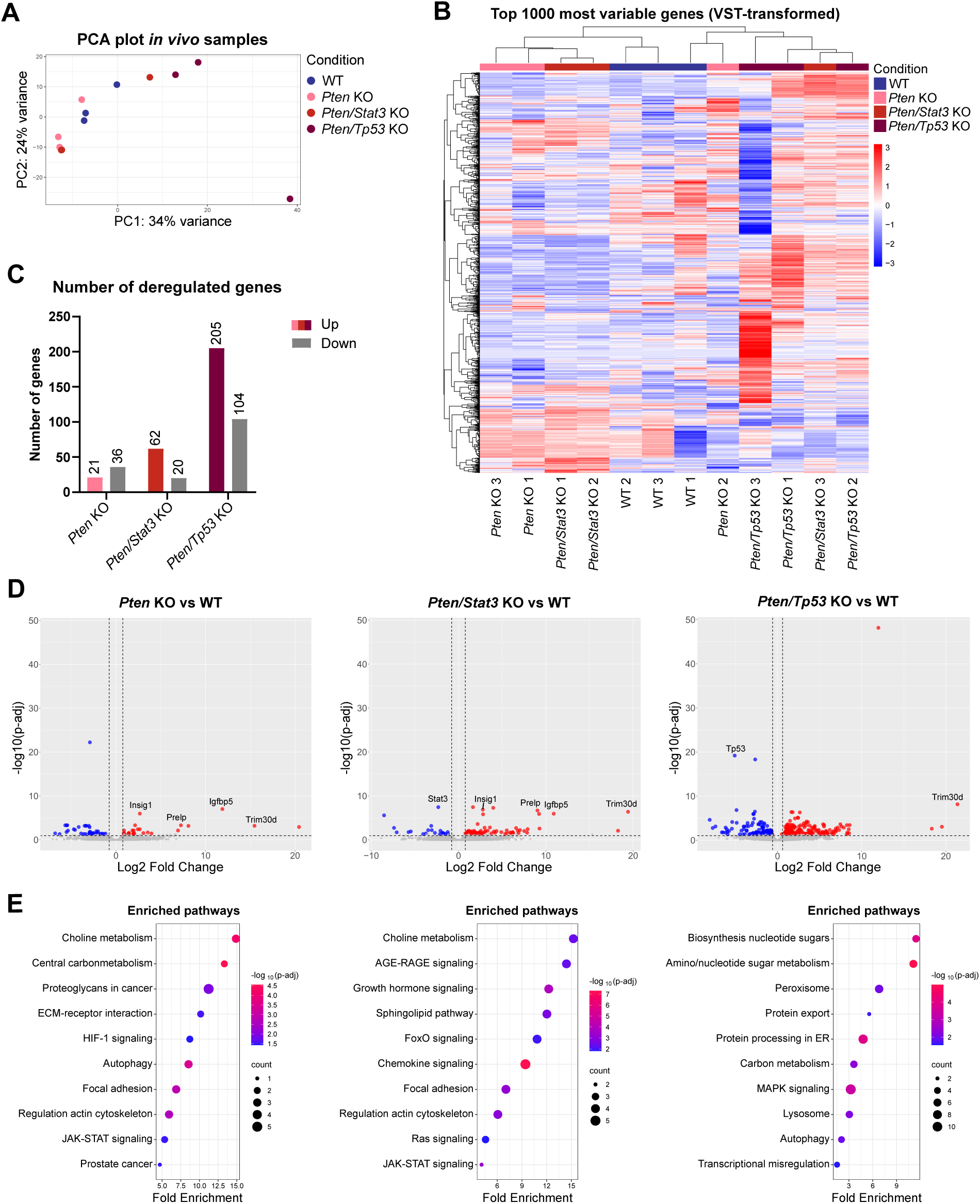
Transcriptomic analysis of PCa tumoroids reveals upregulation of oncogenic signalling and alterations in metabolic pathways. **A**. Principal component analysis (PCA) based on bulk RNA sequencing data from *in vivo* WT organoids and indicated KO tumoroids (biological triplicates per genotype). **B**. Dendrogram and heatmap showing unsupervised hierarchical clustering of the top 1000 most variable genes for all *in vivo* organoids and tumoroids based on VST-normalised gene counts. Rows represent individual genes, while columns represent organoid/tumoroid lines. Colours and intensity reflect expression levels of genes (red: upregulation, blue: downregulation) **C**. Bar graph depicting all significantly overexpressed and downregulated genes (p-adj < 0.05, |Log2fold| > 0) per KO genotype compared to WT organoids (N=3). **D**. Volcano plots depicting differentially expressed genes (DEGs) for *in vivo Pten* KO (left), *Pten/Stat3* KO (middle), and *Pten/Tp53* KO (right) tumoroids compared to WT organoids. Genes with p-adj < 0.05 and Log2fold > 0 (red) or < 0 (blue) are highlighted (N=3). **E**. Bubble plots showing selected significantly enriched pathways based on the KEGG Pathway Database for *in vivo Pten* KO (left), *Pten/Stat3* KO (middle), and *Pten/Tp53* KO (right) tumoroids compared to WT organoids. Size of points reflects number of DEGs mapped to specific pathways, while colour reflects statistical significance (-log10 p-adj) (N=3). See also Figure S2.

Next, we analysed the genotype-specific significant differentially expressed genes (DEGs) between *in vivo* KO tumoroids and WT organoids (Figure 3C,D, Table S5-7). The KO of *Pten* alone, or a dKO of *Pten* and *Stat3*, only resulted in 57 or 82 significant DEGs, respectively. In contrast, combined *Pten* and *Tp53* KO resulted in 309 significant DEGs (Figure 3C). Four genes, which were all reported to interact with the PI3K/AKT pathway, were highly deregulated in all tumoroid genotypes compared to WT organoids (Figure 3D). Among these, the insulin-induced gene 1 (*Insig1*) and the insulin-like growth factor binding protein 5 (*Igfbp5*) are involved in lipid metabolism and can support cell proliferation.^44,45^ Additionally, proline arginine-rich end leucine-rich repeat protein (*Prelp*), an extracellular matrix (ECM) anchoring protein, might be involved in cell adhesion^46^ and epithelial-to-mesenchymal transition (EMT)^47^ (Figure S2F), while tripartite motif-containing 30D (*Trim30d*) is predicted to be a transcription co-activator and possible E3 ubiquitin ligase, and thus might influence several signalling pathways.^48^

To better understand how the identified DEGs impact broader biological processes, we performed KEGG pathway enrichment analysis (Figure 3E) and studied the connections between DEGs using the String database (Figure S3A). Both *Stat3* and *Tp53* appeared as central points in the String networks, validating the dKO tumoroids as representative models to study the changes in protein interactions after genetic deletion of specific genes. In addition, signalling networks of *Prelp*, *Insig1*, and *Igfbp5* were detected. Importantly, in the *Pten* KO and *Pten/Stat3* dKO tumoroids *Pik3r3*, which is part of a regulatory subunit of the PI3K/AKT pathway, showed interactions with integrins, while a network of immune-related proteins was observed in the *Pten/Tp53* dKOs. Among the most significantly enriched KEGG pathways we detected several metabolic pathways including choline metabolism, the central carbon metabolism, the sphingolipid pathway, and amino/nucleotide sugar metabolism, highlighting the influence of *Pten* loss and PI3K/AKT activation on the metabolism of tumoroid lines (Figure 3E).

Along these lines, several PI3K/AKT-dependent signalling pathways such as the JAK/STAT, FOXO, RAS, and MAPK pathways were deregulated in PCa tumoroids of all genotypes. Additionally, in line with the top DEGs we found an enrichment in focal adhesion and regulation of the actin cytoskeleton. The *Pten/Stat3* dKO tumoroids were enriched for chemokine and interferon signalling, which might be a direct effect of the deletion of *Stat3*. While we also detected an enrichment of interferon signalling in the *Pten/Tp53* dKO tumoroids, these tumoroids upregulated pathways involved in protein processing and sugar metabolism indicating increased catabolic needs upon dual loss of *Pten* and *Tp53*. In summary, the loss of *Pten*, *Stat3*, and *Tp53* greatly impacted the transcriptional signatures of tumoroids and highlighted their dependency on PI3K/AKT signalling, which induced the deregulation of major pathways related to metabolism and oncogenic signalling.

### *In vitro* deletion of target genes replicates activation of metabolic pathways and oncogenic signalling observed in *in vivo* KO models

To investigate the effect of target gene deletion on the malignant transformation of healthy organoids, we analysed the differences in gene expression between WT organoids and *in vitro* KO tumoroids, which showed morphological changes upon genetic deletion. For each genotype we analysed three single clones derived from the same maternal line upon tamoxifen induction of the Cre-recombinase. The single clones harbouring either *Pten*, *Pten/Stat3,* or *Pten/Tp53* deletions clustered together based on their genotypes (Figure 4A). Interestingly, the WT control organoids did not group together but clustered in close proximity to the respective KO tumoroid lines derived from the same maternal line. Similarly, hierarchical clustering of the top 1000 most variable genes revealed three clusters, which were dependent on the gene expression of the maternal organoid lines (Figure 4B). Taken together, these results confirmed that the genetic deletion of the target genes *in vitro* changed the gene expression of the organoids but also highlighted the major influence of the transcriptome of their line of origin.

**Figure 4:**
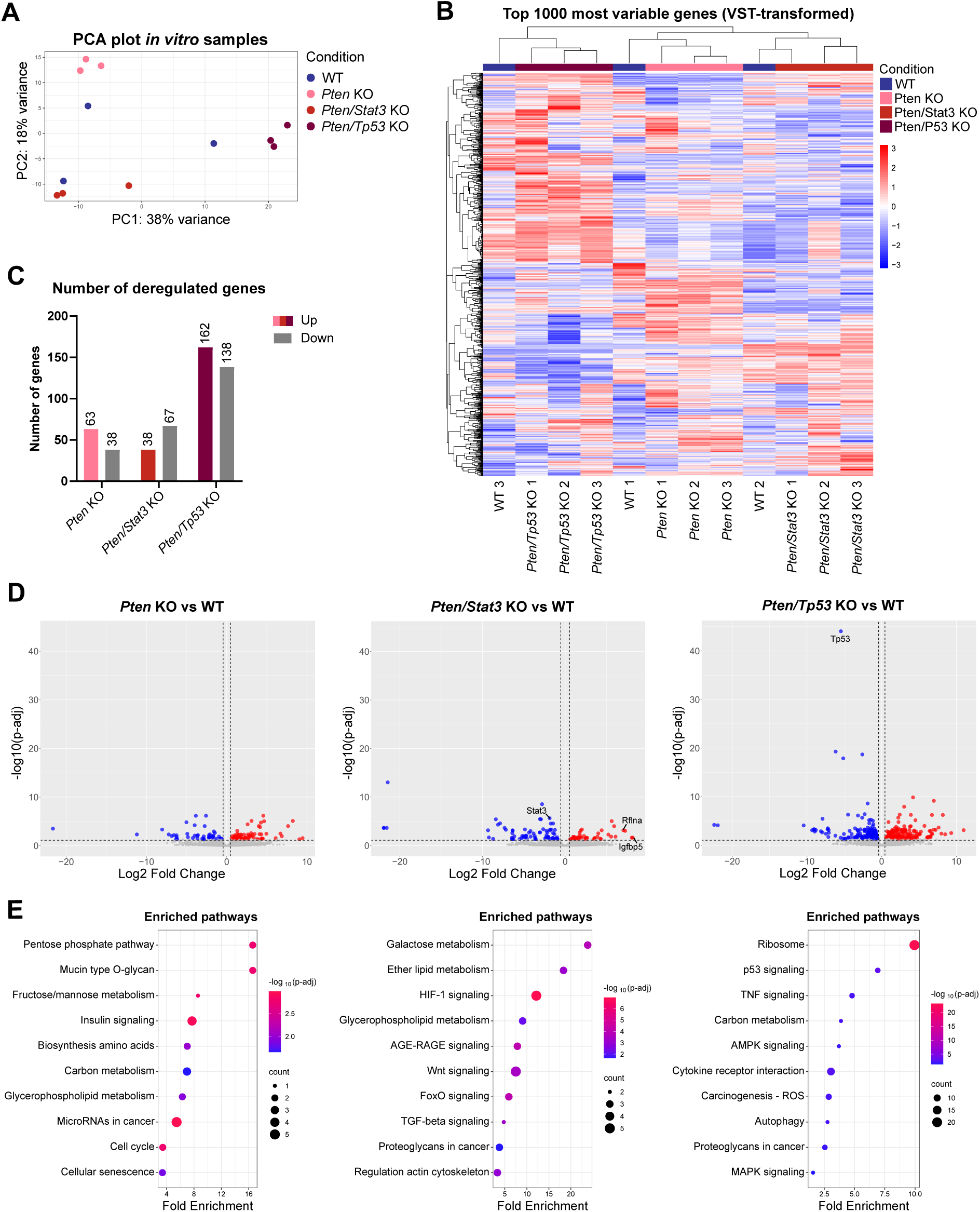
*In vitro* deletion of target genes replicates activation of metabolic pathways and oncogenic signalling observed in *in vivo* KO models. **A**. Principal component analysis (PCA) based on bulk RNA sequencing data of *in vitro* WT organoids and indicated KO tumoroids (single clones/triplicates per genotype). **B**. Dendrogram and heatmap showing unsupervised hierarchical clustering of the top 1000 most variable genes for all *in vitro* organoid and tumoroid lines based on VST-normalised gene counts. Rows represent individual genes, while columns represent organoid/tumoroid lines. Colours and intensity reflect expression levels of genes (red: upregulation, blue: downregulation) **C**. Bar graph depicting all significantly overexpressed and downregulated genes (p-adj < 0.05, |Log2fold| > 0) per KO genotype compared to WT organoids (N=3). **D**. Volcano plots depicting differentially expressed genes (DEGs) for *in vitro Pten* KO (left), *Pten/Stat3* KO (middle), and *Pten/Tp53* KO (right) tumoroids compared to WT organoids. Genes with p-adj < 0.05 and Log2fold > 0 (red) or < 0 (blue) are highlighted (N=3). **E**. Bubble plots showing selected significantly enriched pathways based on the KEGG Pathway Database for *in vitro Pten* KO (left), *Pten/Stat3* KO (middle), and *Pten/Tp53* KO (right) tumoroids compared to WT organoids. Size of points reflects number of DEGs mapped to specific pathways, while colour reflects statistical significance (-log10 p-adj) (N=3).

Next, we focused on the significant DEGs between the *in vitro* KO tumoroids and their WT controls (Figure 4C,D, Table S8-10). The deletion of *Pten* alone, or together with *Stat3* resulted in similar numbers of significant DEGs with 101 or 105 genes, respectively. In line with the *in vivo* 3D cultures, the *Pten/Tp53* dKO tumoroids showed the highest number of DEGs with 300 genes. Out of all DEGs, the phosphofructokinase enzyme (*Pfkm*), a key player in glycolysis^49^, and Refilin A (*Rflna*), which might influence cell adhesion^50^, were significantly upregulated in all *in vitro* tumoroids. Of note, only *Prelp* was overexpressed in all *in vivo* and *in vitro* KO tumoroids.

To identify major deregulated biological processes in the *in vitro* KO tumoroids, we performed pathway enrichment analysis (Figure 4E) and focused on functional String-networks between the DEGs (Figure S3B). *Stat3* and *Tp53* appear as central points in the interaction networks, highlighting that their deletion *in vitro* influences major signalling networks. In addition, the interactions of *Prelp* and *Pfkm* are visible in the networks. Importantly, mimicking the *in vivo Pten/Tp53* dKO tumoroids, an immune-related network of proteins was also observed in the *in vitro Pten/Tp53* dKO models. Similar to the effects observed in *in vivo* tumoroids, pathway enrichment analysis showed major changes in metabolic pathways, including the pentose phosphate pathway, fructose/mannose metabolism, carbon metabolism, and glycerophospholipid metabolism upon deletion of *Pten*, *Pten/Stat3*, or *Pten/Tp53* (Figure 4E). This again highlights the significant role of the PI3K/AKT pathway for metabolic adaptation of cells following *Pten* loss. Interestingly, we found an enrichment in cell cycle and senescence pathways mediated by the upregulation of *Cdkn2a* in the *Pten* KO tumoroids, which has previously been connected to replication stress caused by the loss of tumour suppressor genes.^51^ Similar to the *in vivo* KO tumoroids, we observed the deregulation of major signalling pathways like FOXO, AGE/RAGE, and TGFβ signalling. *In vitro* KO of *Pten* and *Tp53* resulted in deregulation of protein processing as well as autocrine chemokine and cytokine signalling, together promoting cancer-specific processes.

Next, we compared the genotype-specific changes in gene expression between *in vivo* and *in vitro* KO tumoroids. Even though the deletion of *Pten* or the co-deletion of *Pten* and *Stat3* resulted in similar numbers of significant DEGs, the overlap between the *in vivo* and *in vitro* KO tumoroids was around 5%, or 18%, respectively (Figure 5A). The *Pten/Tp53* dKO tumoroids, which showed the highest number of DEGs overall, shared only 16% of all genes between the conditions. Despite this relatively low overlap, gene set enrichment analysis based on the Hallmark Gene Set Collection^52^ revealed an overlap of pathways implicated in tumorigenesis such as the upregulation of KRAS signalling for *Pten* KOs (Figure 5B). In line with the larger overlap of DEGs for the *Pten/Stat3* dKO tumoroids, more hallmark gene sets, including EMT, MTORC1 signalling, and interferon-γ response, were shared between the *in vivo* and *in vitro* tumoroids. Interestingly, seven out of ten hallmark gene sets were identical between the *Pten/Tp53* dKO tumoroids. Of those, different signalling pathways including MTORC1 and P53 signalling, but also pathways influencing the immune response and inflammation, were enriched.

**Figure 5:**
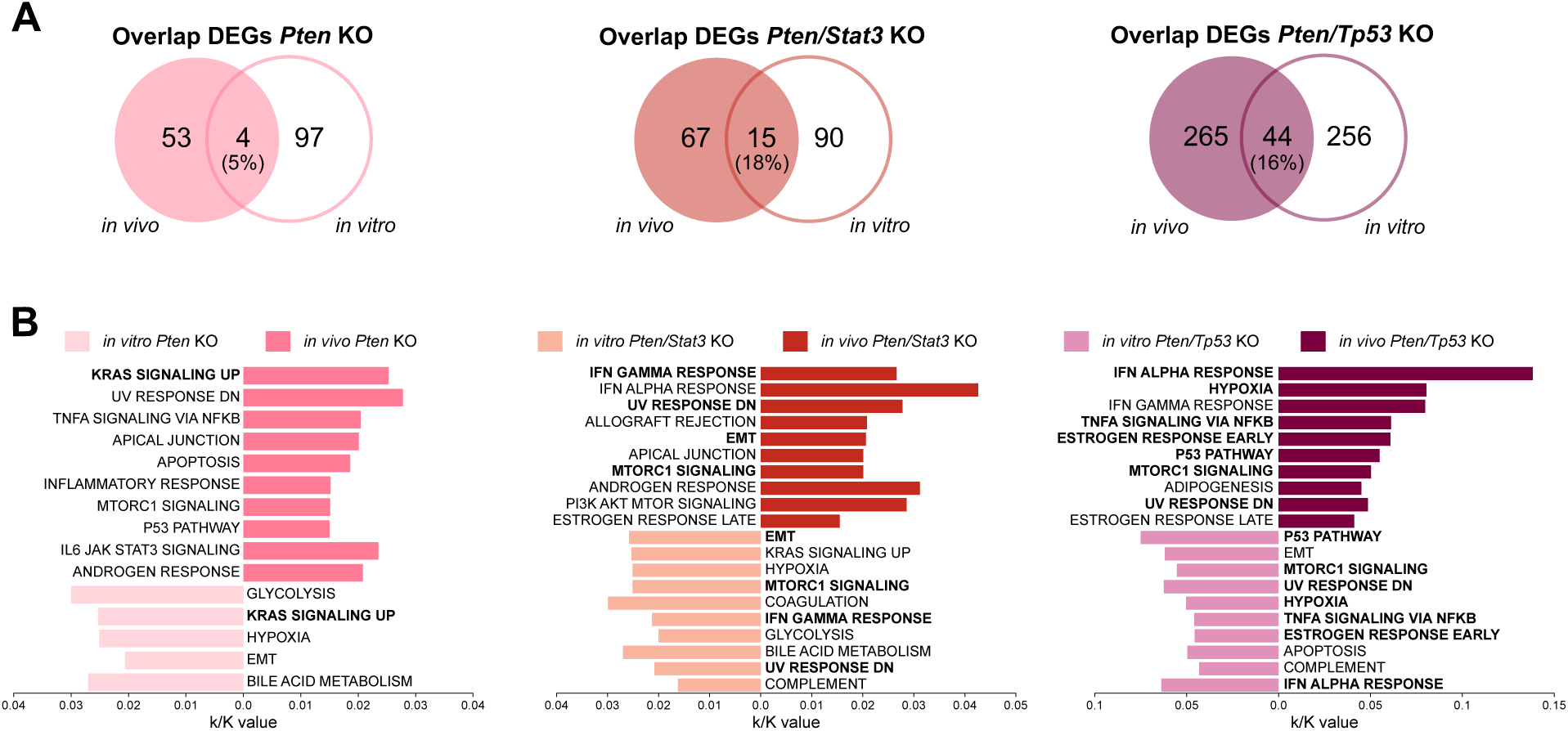
Comparison of differentially expressed genes (DEGs) and enriched hallmark gene sets between *in vivo* and *in vitro* KO tumoroids. **A**. Overlap of significant differentially expressed genes (DEGs) (p-adj < 0.05, |Log2fold| > 0) between *in vivo* and *in vitro Pten* KO (left), *Pten/Stat3* KO (middle), and *Pten/Tp53* KO (right) tumoroids compared to WT organoids (N=3). **B**. Bar graphs depicting the comparison of enriched hallmark gene sets based on MSigDB gene set enrichment analysis of DEGs in (A). k/K value describes the ratio of number of genes in input list (k) divided by the number of total genes in the gene set of the

Of note, the gene sets UV response down and MTORC1 signalling were enriched in all *in vivo* KO tumoroids, indicating a deregulation of stress response, major signalling pathways, and metabolic pathways. The EMT and Hypoxia gene sets were shared between all *in vitro* KO tumoroids hinting to cellular plasticity and an activation of oncogenic pathways after the deletion of the target genes *in vitro*. In conclusion, even though the overlap of significant DEGs is rather small, many biological processes are shared between the *in vivo* and *in vitro* KO tumoroids, especially for the *Pten/Stat3* and *Pten/Tp53* dKO tumoroids. This suggests that major tumour-driving processes can be replicated *in vitro* and depend on cell-intrinsic mechanisms.

### Medium-throughput drug screen identified novel compounds inhibiting PCa tumoroid growth independent of mutational background

As we observed phenotypic and molecular differences between the various KO tumoroid lines, we evaluated the genotype-specific sensitivities of the PCa tumoroids to different pharmaceuticals. For this, we performed a medium-throughput compound screen including 388 common anti-cancer and epigenetic drugs, in addition to selected kinase and pathway inhibitors using the *in vivo* KO tumoroid lines of all three genotypes (Figure 6A, Table S11). For all compounds a percentage of control (POC) value was calculated based on positive and negative controls reflecting 0% and 100% cell viability, respectively.

**Figure 6:**
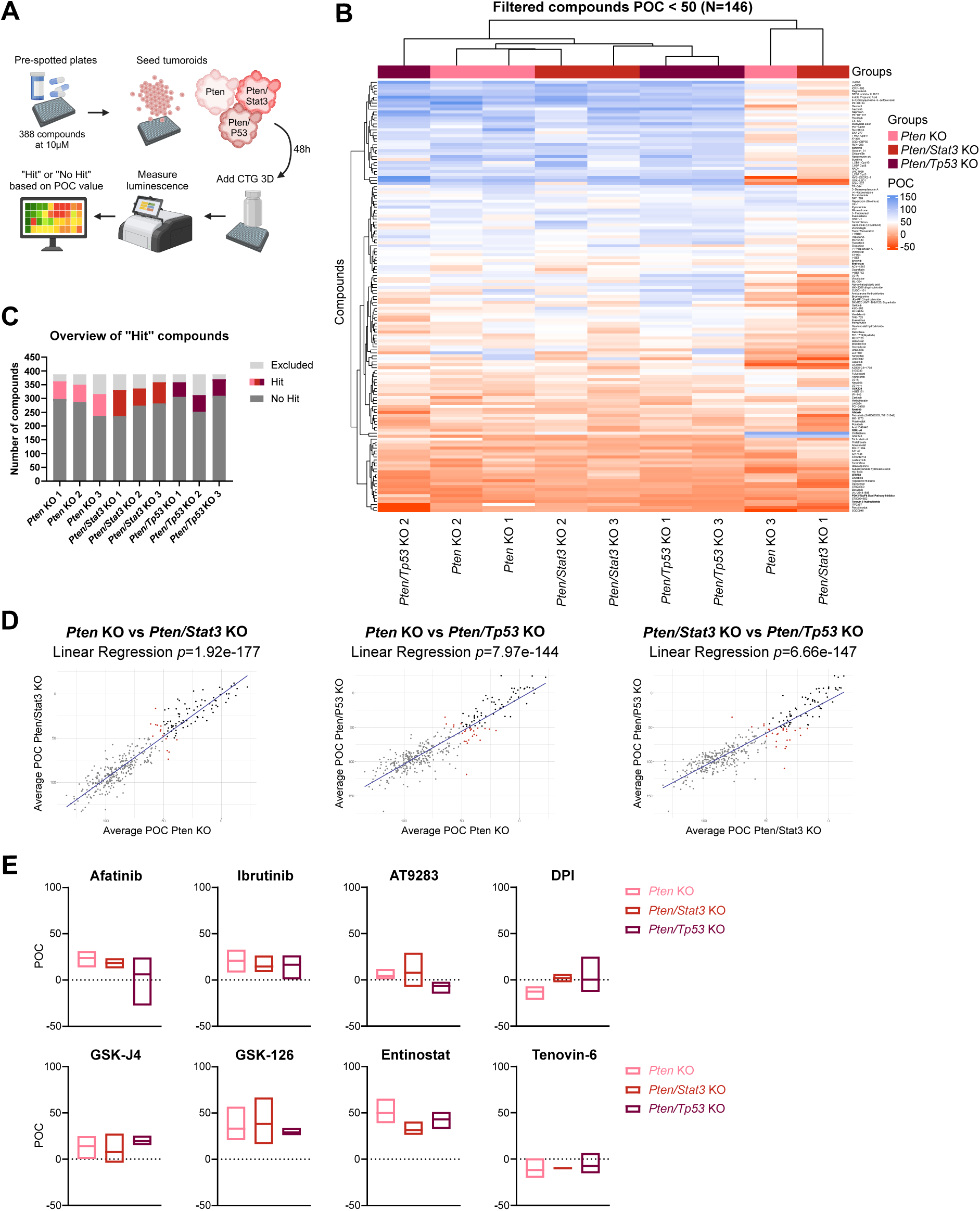
Medium-throughput drug screen identified novel compounds inhibiting PCa tumoroid growth independent of mutational background. **A**. Experimental setup of medium-throughput compound screening. 388 compounds were screened at a single dose of 10µM on small tumoroids with different genetic backgrounds in suspension. CellTiter-Glo® 3D (CTG) was added, and luminescence was measured to calculate the POC (percentage of control) value representing cell viability. **B**. Dendrogram and heatmap showing unsupervised hierarchical clustering of compounds with POC < 50 (N=146) for all tumoroid lines. Rows represent individual compounds, while columns represent *in vivo* KO tumoroids (N=3). Colours and intensity reflect POC values (red: inhibition of growth, blue: no effect). **C**. Bar graph depicting the number of “Hit” (POC < 50) and “No Hit” (POC > 50) compounds per tumoroid line. Compounds were excluded when POC values of duplicates did not match. **D**. Linear regression analysis based on POC values for all compounds for *Pten* KO (N=3) vs *Pten/Stat3* KO (N=3), *p*=1.92e-177 (left); *Pten* KO vs *Pten/Tp53* KO (N=3), *p*=7.97e-144 (middle); and *Pten/Stat3* KO vs *Pten/Tp53* KO, *p*=6.66e-147 (right). Grey: “No Hit”, Black: “Hit”, Red: “Genotype-specific “Hit”. **E**. Box plots showing mean ± SD of POC values for selected compounds effective on all genotypes (N=3 per genotype). Negative POC values are the result of tested compounds showing a higher inhibitory effect than the positive control bortezomib. Statistical analysis was performed using GraphPad Prism 8.0.2 (One-way ANOVA, Tukeýs test). *p* > 0.05 if not specified otherwise. See also Figure S4 and Table S11.

Out of 388 tested compounds, 146 induced POC values lower than 50, representing compounds with distinct anti-cancer effects in the first screening. Hierarchical clustering of these hits showed that two out of three KO lines of each genotype were grouped together, while one line clustered separately (Figure 6B). Interestingly, most compounds effectively inhibited tumoroid growth independent of their mutational background, and the number of Hit-compounds was similar among all lines (Figure 6C). However, for the three tumoroid lines that also did not cluster with their respective replicates in the heatmap, more compounds had to be excluded for further analysis, hinting to technical rather than biological effects. Linear regression analysis of compounds shared between two genotypes further supported the fact that the proliferation of tumoroids with different mutations was effectively inhibited by the same compounds (Figure 6D). Based on these results and previously published literature, we selected eight compounds for further analysis (Figure 6E). Among these were several kinase inhibitors, including the EGFR inhibitor afatinib and the Brutońs kinase inhibitor ibrutinib, but also multitargeted kinase inhibitors AT9283 and the PDPK1/AKT/FLT dual pathway inhibitor (DPI). On the other hand, we focused on four epigenetic modifiers, including the histone demethylase inhibitor GSKJ4, the methyltransferase inhibitor GSK126, and two histone deacetylase (HDAC) inhibitors entinostat and tenovin-6 (T6). Half-maximal inhibitory screening (IC50) revealed that most compounds inhibited tumoroid growth consistently around 1-15µM, while GSKJ4 and GSK126 showed high heterogeneity even between biological replicates. High doses of entinostat, which usually exhibits IC50 values between 0.5 and 10µM on cancer cell lines^53^, were necessary to inhibit PCa tumoroid growth (Figure S4A).

Based on the results of the transcriptomic analysis, which indicated a major role of PI3K/AKT signalling in the metabolic alterations of PCa tumoroids, we chose the DPI, targeting kinases involved in PI3K/AKT signalling, and T6, an inhibitor of sirtuin HDACs and TP53 activator, for further analysis. IC50 screening of *in vivo* and *in vitro* KO tumoroids of all genotypes revealed highly similar sensitivities of both tumoroid models (Figure 7A). Notably, dKO tumoroids showed higher sensitivity to DPI and T6 treatments, with significant differences for *in vivo* dKOs (Figure 7B). Thus, both compounds showed higher efficiencies on more advanced PCa models, and the effect of different genetic deletions on drug response was recapitulated in *in vitro* tumoroids. Of note, the medium-throughput drug screen was performed on tumoroids seeded in suspension, while for the final confirmation all lines were cultured in ECM domes. Even though it has been reported that the ECM can influence the drug response of tumoroids *in vitro*^54^, we did not observe major differences in IC50 values (Figure S4B).

**Figure 7:**
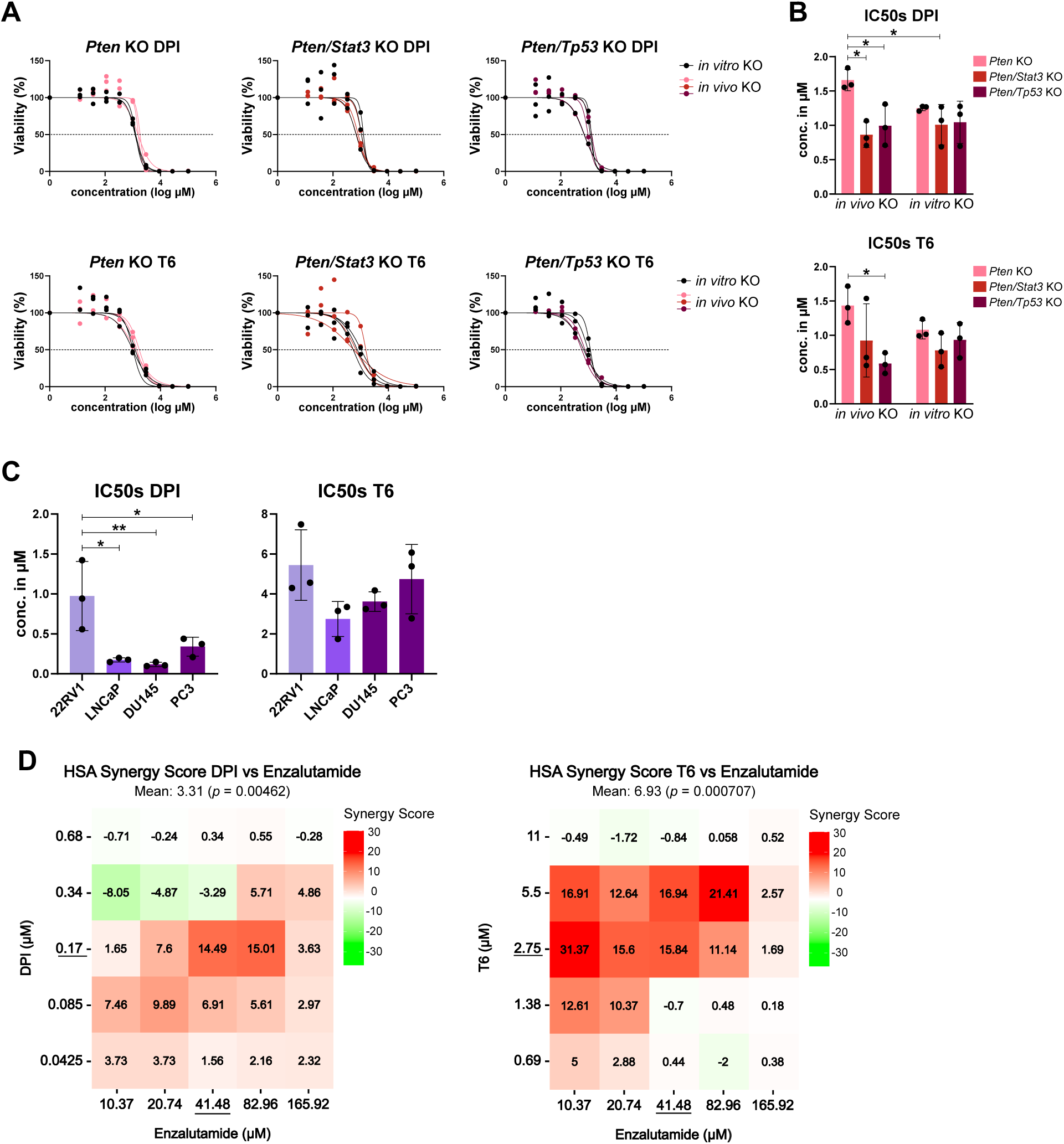
The PDPK1/AKT/FLT dual pathway inhibitor (DPI) and tenovin-6 (T6) show high anti-cancer efficacy in murine tumoroids and human prostate cancer (PCa) cell lines. **A**. Dose-response curves for DPI (top) and T6 (bottom) for *in vivo* and *in vitro Pten* KO (left), *Pten/Stat3* KO (middle), and *Pten/Tp53* KO (right) tumoroids. Points represent means of technical duplicates per tumoroid line (N=3). Curve fitting was performed using GraphPad Prism 8.0.2. **B**. Bar graphs showing means and ± SD of half-maximal inhibitory concentration (IC50) for DPI (top) and T6 (bottom) for *in vivo* and *in vitro* tumoroid lines of all genotypes (N=3). Statistical analysis was performed using GraphPad Prism 8.0.2 (One-way ANOVA, Tukeýs test). *p* > 0.05 if not specified otherwise, \**p* ≤ 0.05. **C**. Bar graphs depicting means and ± SD of IC50 values of DPI (left) and T6 (right) on human PCa cell lines. 22RV1: primary PCa; LNCaP: metastatic PCa; DU145, PC3: metastatic castration-resistant PCa (N=3). Statistical analysis was performed using GraphPad Prism 8.0.2 (One-way ANOVA). *p* > 0.05 if not specified otherwise, \**p* ≤ 0.05; \*\**p* ≤ 0.01. **D**. Heatmaps of synergy scores calculated with the highest single agent (HSA) model for DPI and enzalutamide (left), and T6 and enzalutamide (right) on the human LNCaP cell line. Values > 0 represent synergistic effects, values < 0 represent antagonistic effects. IC50 concentrations of respective compounds are underlined (N=3). See also Figure S4.

To test a potential prognostic significance of genes targeted by DPI and T6 for PCa, we explored the overall survival times of PCa patients dependent on low/high expression levels of the genes of interest.^37^ While the DPI targets, *PDPK1*, *AKT1, AKT2, AKT3,* and *FLT3,* and the T6 targets *SIRT1, SIRT2,* and *DHODH* showed no significant correlation with survival on RNA level, high expression of the T6 target *SIRT3* was significantly associated with worse overall survival (Figure S4C). However, as most of these genes code for effector proteins, their expression on RNA level might not reflect their role for PCa signalling.^55^

To confirm the anti-cancer effect of the selected compounds on human cells, we analysed the cell viability of human PCa cell lines including one primary and three metastatic lines after DPI or T6 treatment (Figure 7C). Indeed, both DPI and T6 effectively inhibited the proliferation of the human cell lines with IC50 concentrations in the same range as for the different tumoroids, confirming that murine PCa tumoroids model the drug response of patient-derived cell line models. Of note, the effect of the two compounds was higher on the metastatic cell lines, indicating that more progressed tumours might be more sensitive to the treatment with DPI and T6. As enzalutamide is one of the most used antiandrogen compounds for the treatment of advanced PCa patients, we used the androgen-responsive LNCaP cell line to investigate whether DPI and T6 could be used in combination with enzalutamide to improve its anti-tumour effect (Figure 7D, Figure S4D). Indeed, both DPI and T6 showed high synergy scores with enzalutamide in the range of the respective IC50 concentrations. Importantly, even low concentrations of enzalutamide in combination with multiple T6 concentrations resulted in high synergy values. In conclusion, both compounds improved the anti-cancer effect of enzalutamide and could thus be beneficial especially for advanced and castration-resistant PCa patients.

## Discussion

In recent years, both human and murine organoid and tumoroid models have been used for studying tumorigenesis and finding novel treatment options for cancer patients as they stably reflect the genetic and epigenetic background, but also the drug response of patients or mouse models.^20^ However, patient-derived prostate organoids and PCa tumoroids have low establishment rates and cannot be maintained for several passages *in vitro*.^25–29^ Here we established a biobank of organoids and PCa tumoroids derived from murine tumours reflecting common patient mutations and compared them to tumoroids generated by genetic deletion of target genes *in vitro*.

Interestingly, RNA sequencing and pathway enrichment analysis of murine PCa tumoroids suggested a metabolic adaption of healthy cells upon deletion of target genes. In humans, both the healthy prostate and PCa tumours exhibit unique metabolic dependencies.^56,57^ While the TCA cycle is suppressed in healthy prostate cells, the development of PCa leads to a metabolic switch by activating the TCA cycle and OXPHOS. Early-stage tumours also heavily rely on lipid and amino acid metabolism for energy production and growth. During advanced and castration-resistant stages of PCa, glycolysis is enhanced (Warburg effect^58^), but OXPHOS and lipid metabolism remain active^56,57^. Importantly, murine PCa models have been used to study PCa metabolism as they recapitulate the metabolic switch observed in humans.^59,60^

Enriched pathways in *in vivo Pten* KO and *Pten/Stat3* dKO tumoroids were mainly mediated pathway.^61^ High *PIK3R3* expression in combination with the loss of *Pten* leads to the constant activation of the PI3K/AKT pathway^62^, which we also observed on protein level. In addition, aberrant lipid metabolism has been observed in PCa, and choline-PET is used to monitor the progression and therapy response of PCa.^63^ We also observed the upregulation of *Pld1*, which mediates PI3K/AKT and mTOR signalling, but also choline metabolism, and thus promotes proliferation and castration resistance in PCa.^64–66^

Even though different genes were deregulated in the *Pten/Tp53* dKO tumoroids, several genes directly involved in glycolysis, including *Hk1*, which is connected to AKT1^67^, *Pgm1*, and *Gfpt1*, were upregulated. Importantly, *Pfkm*, which is also directly involved in glycolysis, showed higher expression in all *in vitro* KO tumoroids. Together, this highlights the major role of the PI3K/AKT pathway in PCa progression but also metabolism of PCa cells.^68^ Importantly, we observed changes in metabolic pathways connected to PCa in tumoroids of all three genotypes. *In vitro* KO tumoroids showed similar enrichment of metabolic signalling upon deletion of target genes, highlighting the potential impact of *Pten* loss on metabolic reprogramming during the first steps of tumorigenesis.

On the other hand, increased activation of the PI3K/AKT pathway together with androgen and TGFβ signalling can induce EMT in PCa to drive metastasis and therapy resistance.^69^ Interestingly, the only gene that was highly upregulated in all *in vivo* and *in vitro* KO tumoroid lines was the proteoglycan *Prelp*, and both its overexpression^47^ and downregulation^70,71^ have been associated with tumour progression and EMT. In colorectal cancer, PRELP interacts with integrins to reduce the stiffness of the ECM to drive metastasis.^47^ As we observed an enrichment of TGFβ signalling and EMT in KO tumoroids, and as the expression of *PRELP* correlates with multiple EMT-related genes in human PCa gene expression data, our results suggest that by remodelling the ECM to increase EMT *Prelp* might lead to PCa progression.

Even though the deletion of *Pten* and the consequent activation of the PI3K/AKT pathway has been explored as a therapeutic option for PCa, most PI3K/AKT inhibitors failed as monotherapies during early clinical testing, mostly due to compensatory signalling mechanisms.^72^ It has been proposed that multitarget kinase inhibitors are a more promising approach for PCa treatment.^73,74^ Out of 388 compounds we identified the DPI^33–35^, which simultaneously inhibits the kinases AKT, and PDPK1 and FLT3 involved in the phosphorylation and thus the complete activation of AKT.^75,76^ DPI has shown promising inhibitory effects in multiple cancer types, notably also in combination with PARP inhibitors.^77–80^ However, although PI3K/AKT signalling plays a major role in PCa development and progression, the inhibitor has not been widely tested as a treatment for PCa.^81^ Here we show that DPI potently inhibits the proliferation of PCa tumoroids and human PCa cell lines, with an even higher effect on more advanced models. As therapy options for advanced PCa patients are limited, DPI could be a novel treatment for these patients.

Apart from metabolic alterations, epigenetic reprogramming is essential for PCa progression and therapy resistance.^82^ Several epigenetic compounds have been investigated as treatment options for PCa, and especially HDAC inhibitors have been tested extensively in preclinical and clinical studies.^83^ Interestingly, our medium-throughput compound screen identified T6 as one of the most effective inhibitors of PCa tumoroid growth. T6 inhibits SIRT1, SIRT2, SIRT3, and the enzyme DHODH.^36,84^ While SIRT1 and SIRT2 can activate the PI3K/AKT pathway to promote proliferation, migration, and neuroendocrine differentiation of PCa cells, SIRT3 usually acts as a tumour suppressor.^85–87^ However, patients with high expression of *SIRT3* in the PRAD-TCGA dataset have shorter overall survival time, hinting to a tumour-promoting effect in PCa. In addition, high *DHODH* expression has been correlated with worse prognosis in PCa patients.^88^ DHODH is involved in the synthesis of pyrimidines, which are needed for the biosynthesis of DNA, RNA, glycoproteins, and phospholipids.^89^ Thus, highly proliferative cancer cells might be more sensitive to inhibition of nucleotide synthesis. Indeed, we observed a stronger effect of T6 on *Pten/Tp53* dKO tumoroids, which showed the highest proliferation rates among the different genotypes. Importantly, we also confirmed the anti-cancer effect of T6 on several human PCa cell lines and propose this compound as a novel therapy option for PCa that has not yet been investigated for this cancer type.

So far, epigenetic compounds are not used as single treatments for solid cancers, and the combination with chemotherapeutics or antiandrogens has shown promising results for PCa.^83^ In line with this, the inhibition of PI3K/AKT in combination with androgen signalling has emerged as a novel treatment strategy.^72^ Importantly, we showed that both DPI and T6 in combination with enzalutamide synergistically inhibit human PCa cell proliferation, and could thus be used to increase the anti-cancer effect of enzalutamide.

Future studies should validate the mechanism of DPI and T6 in *in vitro* and *in vivo* PCa models. Even though tumoroids present useful preclinical models, they do not reflect the complex interactions of cancer cells with the tumour microenvironment.^20^ In addition, we and others^90^ did not observe an effect of the ECM on drug response, but its negative influence has previously been described.^54^ Thus, refining tumoroid culture conditions by incorporating components of the TME and physical stimuli could enhance their physiological relevance for drug development.^20,91^

In conclusion, by using murine PCa tumoroids we identified two promising novel compounds for further validation for PCa treatment. As tumoroids replicated the drug response of human in line with the 3R principles.^92^ Since the deletion of the target genes *in vitro* mirrored the *in vivo* KO tumoroids, the influence of several different mutations on gene expression and drug response could be modelled *in vitro* for a personalised medicine approach.

## Abbreviations

DEGs: differentially expressed genes
DPI PDPK1/AKT/FLT: dual pathway inhibitor
ECM: extracellular matrix
EMT: epithelial to mesenchymal transition
HDAC: histone deacetylase
HE: haematoxylin and eosin
IC50: half-maximal inhibitory concentration
IHC: immunohistochemistry
KO: knockout
PCa: prostate cancer
POC: percentage of control T6 tenovin-6
WT: wildtype

## Resource availability

### Lead contact

Requests for further information and resources should be directed to and will be fulfilled by the lead contact, Gerda Egger.

### Materials availability

All organoid and tumoroid lines generated in this study are available from the lead contact. We are glad to share all models with reasonable compensation by requestor for its processing and shipping, and a completed materials transfer agreement.

### Data and code availability

- Bulk RNA sequencing data was deposited at Gene Expression Omnibus [GSE291912] and are publicly available as of the date of publication.
- This paper analyses existing, publicly available data, accessible at the TCGA (Accession number phs000178).
- No original code has been generated during this study. Publicly available code and packages are cited in the text or method section.
- Any additional information required to reanalyse the data reported in this paper is available from the lead contact upon request.

## Acknowledgements

The authors thank Sabrina Wohlhaupter, Astrid Haase, and Michaela Schlederer for performing IHC stainings, and Martin Raigel for help with pathological analysis. This research was funded in whole by the Austrian Science Fund (FWF) [10.55776/P32771; 10.55776/DOC59; 10.55776/F8300]. ZP was supported by a FFG-FEMtech scholarship (Nr. 8743637). For open access purposes, the author has applied a CC BY public copyright license to any author accepted manuscript version arising from this submission. Figures were partially created using BioRender.

## Author contributions

Conceptualisation: G.E, T.D, and J.K. Methodology: J.K, T.D, Z.P, T.L, T.M, L.V, and S.K. Formal Analysis: J.K, K.D, A.T, G.W. Investigation: J.K, T.D, R.J, E.A, and A.B. Resources: T.L, and L.K. Writing – Original Draft: J.K and G.E. Writing – Review & Editing: all. Visualisation: J.K and G.E. Supervision: G.E. Project Administration: J.K and G.E. Funding Acquisition: G.E and L.K.

## Declaration of Generative AI and AI-assisted technologies in the writing

During the preparation of this work the author used ChatGPT (OpenAI) in order to improve grammar and clarity of the text. After using this tool, the authors reviewed and edited the content as needed and take full responsibility for the content of the publication.

## Methods

### Analysis of human publicly available transcriptomic and genomic data

Data for the Kaplan-Meier survival curves was extracted from the publicly available TCGA PRAD PCa data sets for RNA sequencing and genome/exome sequencing using an online tool.^93^ Survival curves and statistics were performed using GraphPad Prism 8.0.2 (Mantel-Cox test).

### Animal models

All mice were maintained on a C57Bl/6-Sv/129 mixed background under specific pathogen-free conditions at 20-24°C. Previously described PCa mouse models with loxP sites for *Pten*, *Pten/Stat3*, and *Pten/Tp53* were bred with PB-Cre4 mice^94^ to obtain mice with a prostate-specific deletion of respective genes. Tumoroids and organoids were either derived from dKO)^95^, and *Pten*^loxP/loxP^*Trp53*^loxP/loxP^PB-Cre4^+^ (*Pten/Tp53* dKO)^39^ mouse models, or from healthy prostates from *Pten*^loxP/loxP^, *Pten*^loxP/loxP^*Stat3*^loxP/loxP^, or *Pten*^loxP/loxP^*Trp53*^loxP/loxP^ mice (WT), respectively. Male animals of all genotypes were sacrificed at 19 weeks of age and the prostate/tumour tissues were isolated. Only anterior and dorsal lobes were used. Tissue was partly embedded in paraffin or used for organoid/tumoroid generation. All animal experiments were reviewed and approved by the Federal Ministry for Education, Science and Research of the Republic of Austria and conducted according to regulatory and animal well-fare standards (BMWF-66.009/0281-I/3b/2012, BMBWF GZ 66.009/0135-WF/V/3b/2016).

### Organoid and tumoroid establishment

Organoids and tumoroids from murine healthy prostates or prostate tumours, respectively, were isolated and cultured as previously described.^23,43^ Briefly, isolated murine tissues were digested into single cells, the cell pellet was resuspended in Matrigel^R^ (Corning #356234) or Geltrex^TM^ (Gibco #A1413202) and plated as hanging drops. After polymerisation of the matrix, culture medium (Table S1) was added. Organoids and tumoroids were passaged every 5-7 days according to their size and growth rate (0.1% Trypsin, 27G needle). For all experiments organoids/tumoroids below passage 35 were used.

### Lentiviral transduction

For lentiviral transduction, organoid-derived single cells were seeded on 2D tissue culture plates 48h before adding lentiviral particles either generated from the MSCV CreERT2 puro vector (Addgene plasmid # 22776) or the control plasmid MSCV CreCut puro, that was created by shortening the sequence of the Cre-recombinase to make it non-functional. After 48h, transduced cells were plated as 3D cultures in ECM domes, and after 24h selection medium (culture medium + 3.5µg/ml puromycin) was added. Cre-recombinase or CreCut expression was confirmed by PCR (Table S2). After recovery, the KO of the specific genes was induced by adding 500nM 4-hydroxytamoxifen. Final control organoid or KO tumoroid lines were generated from single-cell clones. For this, organoid-derived single cells were seeded sparsely into ECM domes and grown for three days. To dissolve the ECM, Cell Recovery Solution (Corning #354253) was added, and using a microscope and pipette single organoids were transferred to new ECM domes for expansion.

### Western blotting

Protein isolation from snap-frozen organoid/tumoroid pellets was performed as described earlier^96^ with minor changes. For all blots 10µg of protein per sample was used and loaded onto 10% polyacrylamide gels. After incubation with the primary antibodies for the respective genes (Table S3), and incubation with an HRP-conjugated secondary antibody, the signal was developed using chemiluminescent solution ECL (Cytiva Amersham™ ECL™ #RPN2232) and measured using ChemiDoc XRS+ (Bio-Rad). To induce *Tp53* expression, organoids/tumoroids were treated with 100µM CoCl_2_ overnight. Quantification of blots was done using ImageLab software 6.1, and statistical analysis was performed using GraphPad Prism 8.0.2 (One-way ANOVA, Tukeýs test).

### Proliferation assay

Organoid/tumoroid-derived single cells were seeded at a density of 3000 cells/9µl Geltrex^TM^ in a 96-well plate. Medium containing RealTime-Glo (Promega # G9711) was added after ECM polymerisation and refreshed on day three. Luminescence signal was measured every 24h for six days. Data was normalised to signal from day one to calculate the proliferation rate. Statistical analysis was performed using GraphPad Prism 8.0.2 (One-way ANOVA, Tukeýs test).

### Immunohistochemistry

Organoid/tumoroid-derived single cells were grown for seven days at a density of 10,000 cells/15µl Matrigel^R^. After fixation with 4% paraformaldehyde, organoids/tumoroids were resuspended in agarose domes, which were dehydrated and embedded in paraffin. Both murine tissue and embedded 3D lines were cut into 2µm thick sections and stained with haematoxylin and eosin (HE), or with antibodies against proteins of interest (Table S3). Signal was developed using AEC substrate (BD Pharmingen™ #551015) and slides were scanned for further analysis by pathologists trained in uropathology.

### RNA isolation, sequencing, and gene expression analysis

Organoid/tumoroid-derived single cells were seeded at a density of 10,000 cells/15µl Geltrex^TM^. After seven days, ECM domes were dissolved in lysis buffer (Qiagen RNeasy Kit # 74104) and samples were stored at -80°C. RNA was isolated according to the manufactureŕs protocol. The RNA of all generated *in vivo* and *in vitro* organoids and tumoroids was sent to Lexogen GmbH for bulk-RNA sequencing (Illumina shared lane, 100M total reads). Results were mapped to mouse genome GRCm38.101 using STAR aligner and quality control was performed with the RSEQC Quality control package (Python). Changes in gene expression were analysed using DESeq2.^97^ Genes with an adjusted *p*-value < 0.05 and an absolute Log2fold > 0 between groups were considered significant. Data visualisation, including volcano plots and heatmaps, was done using ggplot2.^98^ Biological processes were inferred through pathway enrichment analysis using the pathfindR package.^99^ All analyses were conducted using R version 4.3.1. Gene set enrichment analysis was performed by mapping significantly differentially expressed genes to the mouse-orthologue hallmark gene sets using Mouse MSigDB v2024.1.Mm.^100^ Additional plots were generated using SRPlot.^101^

### Medium-throughput compound screen

All screened compounds were obtained from commercial sources within the PLACEBO in-house collection at the Centre for Molecular Medicine (CeMM) of the Austrian Academy of Sciences, Vienna. Compounds were dispensed into 384-well plates (Corning #3701, #3764) as nanodroplets using an acoustic dispensing system (Echo 550 BeckmanCoulter). *In vivo* tumoroid-derived single cells were grown for two days before isolating small tumoroids using Cell Recovery Solution (Corning #354253). Using an automatic multichannel pipette, 1000 tumoroids were seeded in suspension in 50µl medium supplemented with 5% Geltrex^TM^ per well. For the initial screening, 388 compounds were tested at a concentration of 10µM. As a follow-up, 8-point dose-response curves in a 3-fold dilution series were performed for eight selected compounds following the same protocol. After 48h, viability was measured using CellTiter-Glo 3D Cell Viability Assay (Promega # G9681) and luminescence was measured using the Envision (Revvity) plate reader. DMSO and 10µM bortezomib were used as negative and positive controls, respectively. “Hits” were defined as compounds leading to more than 50% signal inhibition compared to DMSO controls (percentage of control: POC value). Negative POC values are the result of tested compounds showing a higher inhibitory effect than the positive control bortezomib. Initial data analysis of luminescence readouts was performed using Biovia PipelinePilot (Dassault Systems) and Spotfire Analyst (TIBCO) software. Further data analysis and visualisation was performed using the ComplexHeatmap and ggplot2 packages in R. Curve fitting for IC50 calculation and statistical analysis was performed using GraphPad Prism 8.0.2.

### Validation of selected compounds on PCa tumoroids

Dose-response curves for the PDPK1/AKT/FLT dual pathway inhibitor (DPI) (SantaCruz Biotechnology #CAS 331253-86-2) and tenovin-6 (T6) (MedChemExpress #1011301-29-3) were performed on all *in vivo* and *in vitro* tumoroids. Tumoroid-derived single cells were seeded in 9µl Geltrex^TM^ domes at a density of 2000 cells onto 96-well plates and grown for two days, before adding the compounds in a 3-fold dilution series. After 48h, CellTiter-Glo 3D Cell Viability Assay (Promega # G9681) was added to assess cell viability. Half-maximal inhibitory concentration (IC50) was calculated based on negative (max. 0.27% DMSO) and positive (30% DMSO) controls using GraphPad Prism 8.0.2.

### Validation of selected compounds on human PCa cell lines

All PCa cell lines were obtained from ATCC and cultured at 37°C with 5% CO_2_. The human PCa cell lines 22RV1, PC3, and DU145 were cultured in human plasma like medium (HPLM, Gibco #A4899101), while the PCa cell line LNCaP was cultured in RPMI (Gibco #11875085) supplemented with 10% FCS and 1% PenStrep. For IC50 calculations, 4000 cells per cell line were seeded in HPLM per well onto 96-well plates. After cell attachment, the compounds (DPI and T6) or enzalutamide (MedChemExpress #HY-70002) at specified concentrations were added, and Bemcentinib (10 µM) or 30% DMSO were used as positive controls. Cell viability was determined after 48h by adding Resazurin (Sigma Aldrich #B70717) diluted 1:5 in HPLM. After incubation for 2h at 37°C the fluorescence signal was measured (excitation 530/570 nm, emission 580/620nm). Curve fitting and statistical analysis was performed using GraphPad Prism 8.0.2.

### Synergy

For synergy experiments, LNCaP cells were seeded onto 96-well plates at a density of 4000 cells per well in HPLM. After 24h, enzalutamide together with either DPI or T6 were added at specified concentrations. After 48h, cell viability was measured using Resazurin. Data was analysed based on the highest single agent (HSA) synergy model using SynergyFinderplus.^102^

### Statistics

All statistical analysis was performed using GraphPad Prism 8.0.2. Specific statistical tests are mentioned in respective method section and figure legends. 95% confidence interval: ns *p* > 0.05; \**p* ≤ 0.05; \*\**p* ≤ 0.01; \*\*\**p* ≤ 0.001; \*\*\*\**p* ≤ 0.0001.

## Supplemental information

Document S1: Figures S1-4, figure legends for Figures S1-4, supplementary methods, references, Tables S1-4.

Document S2: Table S5-10: DEGs from RNA sequencing (Related to Figure 3, 4)

Document S3: Table S11: List of 388 compounds for medium-throughput compound screen (Related to Figure 6)

## References

1. James, N. D. et al. The Lancet Commission on prostate cancer: planning for the surge in cases. The Lancet 403, 1683–1722 (2024).

2. Crowley, L. & Shen, M. M. Heterogeneity and complexity of the prostate epithelium: New findings from single-cell RNA sequencing studies. Cancer Letters 525, 108–114 (2022).

3. Haffner, M. C. et al. Genomic and phenotypic heterogeneity in prostate cancer. Nat Rev Urol 18, 79–92 (2021).

4. Flores-Téllez, T. del N. J. & Baena, E. Experimental challenges to modeling prostate cancer heterogeneity. Cancer Letters 524, 194–205 (2022).

5. Kaushal, J. B., Takkar, S., Batra, S. K. & Siddiqui, J. A. Diverse landscape of genetically engineered mouse models: Genomic and molecular insights into prostate cancer. Cancer Letters 593, 216954 (2024).

6. Rebello, R. J. et al. Prostate cancer. Nat Rev Dis Primers 7, 1–27 (2021).

7. Almeeri, M. N. E., Awies, M. & Constantinou, C. Prostate Cancer, Pathophysiology and Recent Developments in Management: A Narrative Review. Curr Oncol Rep 26, 1511– 1519 (2024).

8. Yamada, Y. & Beltran, H. The treatment landscape of metastatic prostate cancer. Cancer Letters 519, 20–29 (2021).

9. Cotter, K. & Rubin, M. A. The evolving landscape of prostate cancer somatic mutations. The Prostate 82, S13–S24 (2022).

10. Jamaspishvili, T. et al. Clinical implications of PTEN loss in prostate cancer. Nat Rev Urol 15, 222–234 (2018).

11. Armenia, J. et al. The long tail of oncogenic drivers in prostate cancer. Nat Genet 50, 645–651 (2018).

12. Don-Doncow, N. et al. Expression of STAT3 in Prostate Cancer Metastases. European Urology 71, 313–316 (2017).

13. Abdulghani, J. et al. Stat3 Promotes Metastatic Progression of Prostate Cancer. The American Journal of Pathology 172, 1717–1728 (2008).

14. Han, Z. et al. Inhibition of STAT3 signaling targets both tumor-initiating and differentiated cell populations in prostate cancer. Oncotarget 5, 8416 (2014).

15. Tuo, Z. et al. Pan-cancer analysis of STAT3 indicates its potential prognostic value and correlation with immune cell infiltration in prostate cancer. Discov Onc 15, 654 (2024).

16. Pencik, J. et al. STAT3 regulated ARF expression suppresses prostate cancer metastasis. Nat Commun 6, (2015).

17. Pencik, J. et al. STAT3/LKB1 controls metastatic prostate cancer by regulating mTORC1/CREB pathway. Molecular Cancer 22, 133 (2023).

18. Mai, C.-W. et al. Modeling prostate cancer: What does it take to build an ideal tumor model? Cancer Letters 543, 215794 (2022).

19. Sailer, V. et al. Experimental in vitro, ex vivo and in vivo models in prostate cancer research. Nat Rev Urol 20, 158–178 (2023).

20. Kalla, J., Pfneissl, J., Mair, T., Tran, L. & Egger, G. A systematic review on the culture methods and applications of 3D tumoroids for cancer research and personalized medicine. Cell Oncol. (2024) doi:10.1007/s13402-024-00960-8.

21. Drost, J. et al. Organoid culture systems for prostate epithelial tissue and prostate cancer tissue. Nat Protoc 11, 347–358 (2016).

22. Gao, D. et al. Organoid cultures derived from patients with advanced prostate cancer. Cell 159, 176–187 (2014).

23. Karthaus, W. R. et al. Identification of multipotent luminal progenitor cells in human prostate organoid cultures. Cell 159, 163–175 (2014).

24. Puca, L. et al. Patient derived organoids to model rare prostate cancer phenotypes. Nature Communications 9, (2018).

25. Drost, J. et al. Organoid culture systems for prostate epithelial tissue and prostate cancer tissue. Nat Protoc 11, 347–358 (2016).

26. Brennen, W. N. et al. Defining the challenges and opportunities for using patient-derived models in prostate cancer research. The Prostate 84, 623–635 (2024).

27. Cheaito, K. et al. Establishment and characterization of prostate organoids from treatmentlZlnaïve patients with prostate cancer. Oncology Letters 23, 1–16 (2022).

28. Van Hemelryk, A. et al. Viability Analysis and High-Content Live-Cell Imaging for Drug Testing in Prostate Cancer Xenograft-Derived Organoids. Cells 12, 1377 (2023).

29. Bigot, L. et al. Development of Novel Models of Aggressive Variants of Castration-resistant Prostate Cancer. European Urology Oncology 7, 527–536 (2024).

30. Chan, J. M. et al. Lineage plasticity in prostate cancer depends on JAK/STAT inflammatory signaling. Science 377, 1180–1191 (2022).

31. Gao, X. et al. Blocking PI3K p110β Attenuates Development of PTEN-Deficient Castration-Resistant Prostate Cancer. Molecular Cancer Research 20, 673–685 (2022).

32. Gao, X. et al. Enzalutamide Sensitizes Castration-Resistant Prostate Cancer to Copper-Mediated Cell Death. Advanced Science 11, 2401396 (2024).

33. Zeng, Z. et al. Simultaneous Inhibition of PDK1/AKT and Fms-Like Tyrosine Kinase 3 Signaling by a Small-Molecule KP372-1 Induces Mitochondrial Dysfunction and Apoptosis in Acute Myelogenous Leukemia. Cancer Research 66, 3737–3746 (2006).

34. Koul, D. et al. Inhibition of Akt survival pathway by a small-molecule inhibitor in human glioblastoma. Molecular Cancer Therapeutics 5, 637–644 (2006).

35. Mandal, M. et al. The Akt inhibitor KP372-1 inhibits proliferation and induces apoptosis and anoikis in squamous cell carcinoma of the head and neck. Oral Oncology 42, 430– 439 (2006).

36. Lain, S., et al. Discovery, In Vivo Activity, and Mechanism of Action of a Small-Molecule p53 Activator. Cancer Cell 13, 454–463 (2008).

37. Abeshouse, A. et al. The Molecular Taxonomy of Primary Prostate Cancer. Cell 163, 1011–1025 (2015).

38. Wang, S. et al. Prostate-specific deletion of the murine Pten tumor suppressor gene leads to metastatic prostate cancer. Cancer Cell 4, 209–221 (2003).

39. Chen, Z. et al. Crucial role of p53-dependent cellular senescence in suppression of Pten-deficient tumorigenesis. Nature 436, 725–730 (2005).

40. Limberger, T. et al. KMT2C methyltransferase domain regulated INK4A expression suppresses prostate cancer metastasis. Mol Cancer 21, 89 (2022).

41. Lee, M., Kang, H. & Jang, S.-W. CoCl2 induces PC12 cells apoptosis through p53 stability and regulating UNC5B. Brain Research Bulletin 96, 19–27 (2013).

42. Choudhury, A. D. PTEN-PI3K pathway alterations in advanced prostate cancer and clinical implications. The Prostate 82, S60–S72 (2022).

43. Chua, C. W. et al. Single luminal epithelial progenitors can generate prostate organoids in culture. Nat Cell Biol 16, 951–4 (2014).

44. Waters, J. A., Urbano, I., Robinson, M. & House, C. D. Insulin-like growth factor binding protein 5: Diverse roles in cancer. Front. Oncol. 12, (2022).

45. Ouyang, S., Mo, Z., Sun, S., Yin, K. & Lv, Y. Emerging role of Insig-1 in lipid metabolism and lipid disorders. Clinica Chimica Acta 508, 206–212 (2020).

46. Li, X. et al. PRELP inhibits colorectal cancer progression by suppressing epithelial-mesenchymal transition and angiogenesis via the inactivation of the FGF1/PI3K/AKT pathway. Apoptosis 30, 16–34 (2025).

47. Gui, Y., Deng, X., Li, N. & Zhao, L. PRELP reduce cell stiffness and adhesion to promote the growth and metastasis of colorectal cancer cells by binding to integrin α5. Experimental Cell Research 441, 114151 (2024).

48. Offermann, A. et al. Analysis of tripartite motif (TRIM) family gene expression in prostate cancer bone metastases. Carcinogenesis 42, 1475–1484 (2021).

49. Gao, W. et al. The role of S-nitrosylation of PFKM in regulation of glycolysis in ovarian cancer cells. Cell Death Dis 12, 1–14 (2021).

50. Shao, F. et al. Establishing a metastasis-related diagnosis and prognosis model for lung adenocarcinoma through CRISPR library and TCGA database. J Cancer Res Clin Oncol 149, 885–899 (2023).

51. Jung, S. H. et al. mTOR kinase leads to PTEN-loss-induced cellular senescence by phosphorylating p53. Oncogene 38, 1639–1650 (2019).

52. Liberzon, A. et al. Molecular signatures database (MSigDB) 3.0. Bioinformatics 27, 1739–1740 (2011).

53. Entinostat (MS-275) | HDAC Class I Inhibitor | MedChemExpress. MedchemExpress.com https://www.medchemexpress.com/Entinostat.html.

54. Jung, D. J. et al. A one-stop microfluidic-based lung cancer organoid culture platform for testing drug sensitivity. Lab Chip 19, 2854–2865 (2019).

55. Buccitelli, C. & Selbach, M. mRNAs, proteins and the emerging principles of gene expression control. Nat Rev Genet 21, 630–644 (2020).

56. Ahmad, F., Cherukuri, M. K. & Choyke, P. L. Metabolic reprogramming in prostate cancer. Br J Cancer 125, 1185–1196 (2021).

57. Pujana-Vaquerizo, M., Bozal-Basterra, L. & Carracedo, A. Metabolic adaptations in prostate cancer. Br J Cancer 131, 1250–1262 (2024).

58. DeBerardinis, R. J. & Chandel, N. S. We need to talk about the Warburg effect. Nat Metab 2, 127–129 (2020).

59. Wang, L. et al. PTEN-knockout regulates metabolic rewiring and epigenetic reprogramming in prostate cancer and chemoprevention by triterpenoid ursolic acid. The FASEB Journal 36, e22626 (2022).

60. Sun, J. et al. Resistance to Androgen Deprivation Leads to Altered Metabolism in Human and Murine Prostate Cancer Cell and Tumor Models. Metabolites 11, 139 (2021).

61. Lin, W. et al. PIK3R3 is upregulated in liver cancer and activates Akt signaling to control cancer growth by regulation of CDKN1C and SMC1A. Cancer Medicine 12, 14413– 14425 (2023).

62. Xu, W., Yu, M., Qin, J., Luo, Y. & Zhong, M. LACTB Regulates PIK3R3 to Promote Autophagy and Inhibit EMT and Proliferation Through the PI3K/AKT/mTOR Signaling Pathway in Colorectal Cancer. Cancer Management and Research 12, 5181–5200 (2020).

63. Zhou, S. et al. Value of 11C-Choline PET/CT-Based Multi-Metabolic Parameter Combination in Distinguishing Early-Stage Prostate Cancer From Benign Prostate Diseases. Front. Oncol. 10, (2021).

64. Zhou, W. et al. Inhibition of Phospholipase D1 mRNA Expression Slows Down the Proliferation Rate of Prostate Cancer Cells That Have Transited to Androgen Independence. J Cancer 9, 3620–3625 (2018).

65. Borel, M., Cuvillier, O., Magne, D., Mebarek, S. & Brizuela, L. Increased phospholipase D activity contributes to tumorigenesis in prostate cancer cell models. Mol Cell Biochem 473, 263–279 (2020).

66. Utter, M. et al. Elevated phospholipase D activity in androgen-insensitive prostate cancer cells promotes both survival and metastatic phenotypes. Cancer Letters 423, 28–35 (2018).

67. Yu, Y. et al. AKT1 Promotes Tumorigenesis and Metastasis by Directly Phosphorylating Hexokinases. Journal of Cellular Biochemistry 125, e30613 (2024).

68. Hoxhaj, G. & Manning, B. D. The PI3K-AKT network at the interface of oncogenic signalling and cancer metabolism. Nat Rev Cancer 20, 74–88 (2020).

69. Papanikolaou, S., Vourda, A., Syggelos, S. & Gyftopoulos, K. Cell Plasticity and Prostate Cancer: The Role of Epithelial–Mesenchymal Transition in Tumor Progression, Invasion, Metastasis and Cancer Therapy Resistance. Cancers 13, 2795 (2021).

70. Kosuge, H. et al. Proteomic identification and validation of novel interactions of the putative tumor suppressor PRELP with membrane proteins including IGFI-R and p75NTR. Journal of Biological Chemistry 296, (2021).

71. Hong, R. et al. PRELP has prognostic value and regulates cell proliferation and migration in hepatocellular carcinoma. J Cancer 11, 6376–6389 (2020).

72. Braglia, L., Zavatti, M., Vinceti, M., Martelli, A. M. & Marmiroli, S. Deregulated PTEN/PI3K/AKT/mTOR signaling in prostate cancer: Still a potential druggable target? Biochimica et Biophysica Acta (BBA) - Molecular Cell Research 1867, 118731 (2020).

73. Imamura, J. et al. Lineage plasticity and treatment resistance in prostate cancer: the intersection of genetics, epigenetics, and evolution. Front. Endocrinol. 14, (2023).

74. Bello, T. et al. Computational modeling identifies multitargeted kinase inhibitors as effective therapies for metastatic, castration-resistant prostate cancer. Proc Natl Acad Sci U S A 118, e2103623118 (2021).

75. Zheng, N., Wei, J., Wu, D., Xu, Y. & Guo, J. Master kinase PDK1 in tumorigenesis. Biochimica et Biophysica Acta (BBA) - Reviews on Cancer 1878, 188971 (2023).

76. Takahashi, S. Downstream molecular pathways of FLT3 in the pathogenesis of acute myeloid leukemia: biology and therapeutic implications. J Hematol Oncol 4, 13 (2011).

77. Zhao, Y. et al. SoNar, a Highly Responsive NAD+/NADH Sensor, Allows High-Throughput Metabolic Screening of Anti-tumor Agents. Cell Metabolism 21, 777–789 (2015).

78. Viera, T. & Patidar, P. L. DNA damage induced by KP372-1 hyperactivates PARP1 and enhances lethality of pancreatic cancer cells with PARP inhibition. Sci Rep 10, 20210 (2020).

79. Jiang, L. et al. KP372-1-Induced AKT Hyperactivation Blocks DNA Repair to Synergize With PARP Inhibitor Rucaparib via Inhibiting FOXO3a/GADD45α Pathway. Front. Oncol. 12, (2022).

80. Jiao, B. et al. Bladder cancer selective chemotherapy with potent NQO1 substrate co-loaded prodrug nanoparticles. Journal of Controlled Release 347, 632–648 (2022).

81. Guo, J. et al. Differential Sensitization of Different Prostate Cancer Cells to Apoptosis. Genes Cancer 1, 836–846 (2010).

82. Macedo-Silva, C. et al. Epigenetic mechanisms underlying prostate cancer radioresistance. Clin Epigenet 13, 125 (2021).

83. Kumaraswamy, A. et al. Recent Advances in Epigenetic Biomarkers and Epigenetic Targeting in Prostate Cancer. European Urology 80, 71–81 (2021).

84. Ladds, M. J. G. W. et al. Exploitation of dihydroorotate dehydrogenase (DHODH) and p53 activation as therapeutic targets: A case study in polypharmacology. J Biol Chem 295, 17935–17949 (2021).

85. Chouhan, S., Muhammad, N., Usmani, D., Khan, T. H. & Kumar, A. Molecular Sentinels: Unveiling the Role of Sirtuins in Prostate Cancer Progression. International Journal of Molecular Sciences 26, 183 (2025).

86. Cui, Y. et al. Effect of SIRT1 Gene on Epithelial-Mesenchymal Transition of Human Prostate Cancer PC-3 Cells. Med Sci Monit 22, 380–386 (2016).

87. Ruan, L., Wang, L., Wang, X., He, M. & Yao, X. SIRT1 contributes to neuroendocrine differentiation of prostate cancer. Oncotarget 9, 2002–2016 (2017).

88. Guo, S. et al. DHODH inhibition represents a therapeutic strategy and improves abiraterone treatment in castration-resistant prostate cancer. Oncogene 43, 1399–1410 (2024).

89. Zhou, Y. et al. DHODH and cancer: promising prospects to be explored. Cancer Metab 9, 22 (2021).

90. Lumibao, J. C. et al. The effect of extracellular matrix on the precision medicine utility of pancreatic cancer patient–derived organoids. JCI Insight 9, e172419.

91. Liu, H., Gan, Z., Qin, X., Wang, Y. & Qin, J. Advances in Microfluidic Technologies in Organoid Research. Advanced Healthcare Materials 13, 2302686 (2024).

92. Kiani, A. K., et al. Ethical considerations regarding animal experimentation. J Prev Med Hyg 63, E255–E266 (2022).

93. Smith, J. C. & Sheltzer, J. M. Genome-wide identification and analysis of prognostic features in human cancers. Cell Reports 38, 110569 (2022).

94. Wu, X. et al. Generation of a prostate epithelial cell-specific Cre transgenic mouse model for tissue-specific gene ablation. Mech Dev 101, 61–69 (2001).

95. Pencik, J. et al. STAT3 regulated ARF expression suppresses prostate cancer metastasis. Nature Communications 6, 7736 (2015).

96. Zrimšek, M. et al. Quantitative Acetylomics Uncover Acetylation-Mediated Pathway Changes Following Histone Deacetylase Inhibition in Anaplastic Large Cell Lymphoma. Cells 11, 2380 (2022).

97. Love, M. I., Huber, W. & Anders, S. Moderated estimation of fold change and dispersion for RNA-seq data with DESeq2. Genome Biology 15, 550 (2014).

98. Wickham, H. Ggplot2. (Springer International Publishing, Cham, 2016). doi:10.1007/978-3-319-24277-4.

99. Ulgen, E., Ozisik, O. & Sezerman, O. U. pathfindR: An R Package for Comprehensive Identification of Enriched Pathways in Omics Data Through Active Subnetworks. Front Genet 10, 858 (2019).

100. Subramanian, A. et al. Gene set enrichment analysis: A knowledge-based approach for interpreting genome-wide expression profiles. Proceedings of the National Academy of Sciences 102, 15545–15550 (2005).

101. Tang, D., et al. SRplot: A free online platform for data visualization and graphing. PLOS ONE 18, e0294236 (2023).

102. Zheng, S. et al. SynergyFinder Plus: Toward Better Interpretation and Annotation of Drug Combination Screening Datasets. Genomics Proteomics Bioinformatics 20, 587– 596 (2022).

